# Shock or jump: deimatic behaviour is repeatable and polymorphic in a yellow-bellied toad

**DOI:** 10.1101/2022.04.29.489992

**Authors:** Andrea Chiocchio, Giuseppe Martino, Roberta Bisconti, Claudio Carere, Daniele Canestrelli

## Abstract

Inter-individual variation in antipredatory strategies has long attracted curiosity among scientists. Deimatisms is a complex and time-structured antipredatory strategy consisting in prey suddenly unleashing unexpected defences to frighten predators and stop their attack. Being deimatism traditionally considered as a stereotyped antipredatory response, the inter-individual variation in phenotypic traits related to deimatic displays is almost unexplored. In this study, we employed common garden experiments on 71 yellow-bellied toad *Bombina pachypus* to investigate the extent and pattern of inter-individual variation in the *unken-reflex* behaviour, a deimatic display performed by some amphibians. Results show that deimatic displays consistently differ among individuals. Only about half of the individuals reacted to the predation stimuli by exhibiting the display, which varied in responsiveness, duration and intensity. All the investigated descriptors were repeatable (R > 0.50, p < 0.01). Finally, we found significant correlations between the measured parameters, defining two alternative behavioural profiles: individuals quickly doing *unken-reflex*, with high intensity and long duration of the display, and individuals avoiding *unken-reflex* but rather escaping. Such dichotomy resembles respectively the proactive and reactive coping styles. Such an unexpected variation in deimatic behaviour raises intriguing questions on the evolutionary processes shaping multiple adaptive responses to predation within populations.

## Introduction

The boundless variety of antipredatory strategies spread in the animal kingdom has long attracted curiosity among scientists and inspired evolutionary ecology research through the last two centuries [1]. Antipredatory strategies typically involve the concerted evolution of suites of visual, chemical, acoustic, and behavioural traits, which allow the prey to survive the encounter with the predator [2,3,4,5,6,7]. Several studies reported large within-species variation in phenotypic traits related to the antipredatory strategies. Interestingly, a large part of this variation regards the individual ability to actively modulate the antipredatory responses. Still, inter-individual variation has been reported also in more complex anti-predator strategies where variation was traditionally considered as non-adaptive, such as camouflage and aposematism [8,9]. Such variation has been explained by differential selection exerted by predators on prey populations [10,11,12,13,14,15,16], and by the interaction between contrasting evolutionary forces and ecological trade-offs acting on the many phenotypic traits involved (e.g. balancing the costs of foraging and mating with the costs of sequestering/synthesizing compounds for colorful signals and/or chemical defenses [17,18,19]. However, the data available do not allow a full understanding of the processes maintaining this variation within populations.

Deimatism is an appealing antipredatory strategy, which consists in displaying striking and transitory behavioral responses aimed at inhibiting predator attacks (i.e. startling behaviours), and it is spread among several taxa, especially arthropods, cephalopods, and amphibians [1,9,20,21,22,23]. Deimatic displays are multicomponent as they combine multiple defensive strategies, such as camouflage and aposematism, and typically involve either chromatic and behavioural components [23,24]. Furthermore, deimatic displays are dynamic behaviors as individuals can actively modulate duration, frequency, and intensity of the exhibition depending e.g. on the type and/or behaviour of predators and the prey risk perception. Thus, deimatic species provide an appealing system to investigate the trade-offs between alternative behavioural phenotypes when facing with a threat. Yet, the extent of inter-individual variation in deimatic behavior within and among populations is almost unexplored for the most of deimatic species.

In this study, we investigate the extent of inter-individual variation in deimatic behaviour in the Apennine yellow-bellied toad *Bombina pachypus. Bombina* toads are textbook examples of the *unken-reflex*, a deimatic behavior exhibited by many other toads and salamanders. These species usually bear cryptic colorations on their dorsal side. However, when the attack by a predator is perceived as imminent and unavoidable, these toads contract dorsal muscles, suddenly arch their body, expose their vividly colored ventral side, and release toxins from their skin glands as a reinforcement to the warning signal [25,26]. Preliminary evidence showed some level of variation in the *unken-reflex* behaviour among distinct populations in either *B. variegata* and *B. bombina* [27,28]. Such variation has been linked to the presence of different predator sets among the tested populations [28]. However, there are no data on the extent of this variation among individuals within populations, which cannot be imputed to differential predator selective pressures.

Here, we collected Apennine yellow-bellied toads from southern Italy and scored their *unken-reflex* behaviour in a series of common garden experiments, with the aim to address the following questions: (i) Is the *unken reflex* polymorphic and repeatable within populations? (ii) What are the main components of its variation?

## Material and Methods

A total of 71 *B. pachypus* male individuals were collected during the spring 2020 from a number of close ponds in the surroundings of Reggio Calabria (Aspromonte massif, southern Italy: 38°00’N, 15°51’E). We collected adult males from close but distinct ponds to ensure similar environmental conditions and predators, and to avoid collecting related individuals. Also, we selected this population because it has not been affected by either recent and historical decline, and all the individuals can be considered belonging to a unique and panmittic population [29]. Details about the study species, sampling and housing procedures are provided as Supplementary material.

### Behavioural essays

To obtain behavioural descriptors of the *unken-reflex* response under predator attack, *B. pachypus* individual were submitted to the Tonic Immobility (TI) test, a standardized method that induces thanatosis-like antipredatory reactions in many species of different taxa, including amphibians [30,31]. The TI test consists in simulating the attack of a predator by hitting or catching the prey head for few seconds, and then scoring qualitative and quantitative parameters describing the prey immobility reaction. In the case of *unken-reflex*, toads under threat are expected to arch their body and expose their vividly colored ventral side for the time necessary to the predator to recognize the aposematic signal and give up the attack. Thus, we (i) simulated the attack of a predator in artificial condition, (ii) recorded the toad reaction by means of a camera, and (iii) scored the parameters describing the *unken-reflex* reaction, i.e. latency to react, duration and intensity.

Before the simulation, toads were individually placed in the centre of a white container (50 × 30 × 10 cm) and covered with a white plastic cup for two minutes. Then, the plastic cup was raised up and the stimulation began. The stimulation consisted in gentle hitting the head of the toad with a finger for a maximum of 60 seconds. If a toad moved away, the stimulation continued for a maximum of 60 seconds, then it was considered that *unken-reflex* had not been induced and a score of 0 s was given for all the parameters. Conversely, if toads arched the body, not showing any movement, TI immobility parameters were measured. If the *unken-reflex* lasted for more than 5 minutes, the test was terminated and a score of 300 s was given for duration. Animals were always handled by the same researcher (GM). The tests were recorded with a camera (Panasonic FZ300) fixed at 50 cm from the container, and the recordings were evaluated by the same operator, who scored all the response variables. The following variables were measured to characterize the *unken-reflex* reaction: *unken-reflex* occurrence, expressed as *yes* or *not*; latency, expressed as the time delay in showing the *unken-reflex* after the test began (from 0 to 60); the duration time of the immobility, in seconds (from 0 to 300); the intensity of the response, expressed as a scale from 0 to 3 where 0 is no response, 1 is light response (only the feet are lift up), 2 is moderate response (all the limbs are lift up), and 3 is strongest reaction (the limbs are lift up and the whole body is arched). All the variables were measured using the Boris 7.9.6 software package for video analysis [32]. The test was repeated for each individual under the same conditions, after a seven days interval.

### Statistical analyses

Individual behavioral consistency was assessed by calculating repeatability estimates on repeated measures of each measured parameter (i.e. latency, duration and intensity of the reaction) using mixed models [33]. Repeatability coefficients R, that is, intraclass correlation coefficients, for each parameter were calculated using the “rpt.” function of the “rptR” package [34,35] in R software (version 4.0.0 [36]). Repeatability was estimated as the ratio of between-individual variance to total variance with generalized linear mixed effects models (GLMM) for Poisson distributed data, using individual identity as a random factor and the link-scale approximation of R [33]; for assessing repeatability for *unken-reflex* occurrence, we applied the “rpt.” function for binary data. The 95% confidence intervals (CI) around repeatability estimates were generated by performing 1,000 parametric bootstrap iterations. Variables were considered ‘highly’ repeatable if R > 0.5 or ‘marginally’ repeatable if R > 0.2, and the lower bound of the CI was >0.0 [37,38,39].

Correlation among the measured behavioural descriptors was tested using the Spearman’s rank correlation method implemented in the “spearman.test” function of the “pspearman” R package [40].

## Results

**Results** from the TI tests showed inter-individual variation in the *unken-reflex* deimatic behaviour within the investigated *B. pachypus* population, as well as consistency in the individual response across trials. During the first trial, 38 out of 71 individuals (53.5%) reacted to the predation stimuli performing *unken-reflex*; all the other individuals tried to escape without showing any deimatic display. During the second trial, 41 out of 71 individuals (57.7%) performed the *unken-reflex* display. The *unken-reflex* response was highly repeatable (R = 0.56, CI= 0.16 - 0.70, P = 7.34E-05).

We also found marked inter-individual variation in the way individual performed *unken-reflex*. Latency spanned from 0.24 to 42.59 s in the first trial (mean 8.83 s, median 3.88, sd.± 11.73) and from 0.40 s to 25 s in the second trial (mean 5.77 s, median 4.18, sd.± 6.22); duration spanned from 0.31 to 236.50 s in the first trial (mean 56.11 s, median 22.03s, sd.± 68.03) and from 0.01 to 157.99 s in the second trial (mean 38.88 s, median 12.47, sd.± 49.69); intensity spanned from 1 to 3 in the first trial (mean 2.37, median 3, sd.± 0.79s) and from 1 to 3 in the second trial (mean 2.22, median 3, sd.± 0.88). We found consistency in the individual behavior between the two trials: all the measured behavioural descriptors were significantly and highly repeatable (Table 1). Finally, they were significantly correlated: fast responders showed high intensity and long duration of the deimatic behaviour (Table 2).

**Table 1.** Repeatability estimates of the *unken-reflex* behavioural descriptors among different trials, obtained by fitting the generalized linear mixed effects models (GLMM) implemented in the “rptR” R package. R: repeatability; SE, standard error; CI, 95% confidence intervals obtained from 1,000 parametric bootstrap iterations; P: significance, obtained by likelihood ratio tests. N = 71.

|  | Repeatability |  |  |  |
| --- | --- | --- | --- | --- |
|  | Occurrence | Intensity | Latency | Duration |
| **R** | 0.56 | 0.65 | 0.70 | 0.80 |
| **SE** | 0.15 | 0.08 | 0.07 | 0.07 |
| **CI** | [0.16, 0.70] | [0.47, 0.77] | [0.55, 0.82] | [0.64, 0.90] |
| **P [LRT]** | 7.34E-05 | 1.04E-08 | 2.22E-08 | 8.68E-11 |

**Table 2.** Matrix of the Spearman’s rank correlation coefficients (below) and significance (above) among the 3 descriptors of *unken-reflex* reaction (latency, intensity and duration). N = 71.

|  | Spearman’s Rho |  |  |
| --- | --- | --- | --- |
|  | Latency | Intensity | Duration |
| **Latency** | - | 0.033 | 0.046 |
| **Intensity** | -0.35 | - | 0.004 |
| **Duration** | -0.32 | 0.46 | - |

Results of the first trial are shown in Figure 1 and highlight two alternative behavioral strategies: about half of the individuals did not perform *unken-reflex* (latency = 60 s, intensity and duration = 0); on the other hand, most of the individuals performing *unken-reflex* showed low latency, high intensity and long duration of the reaction.

**Figure 1.**
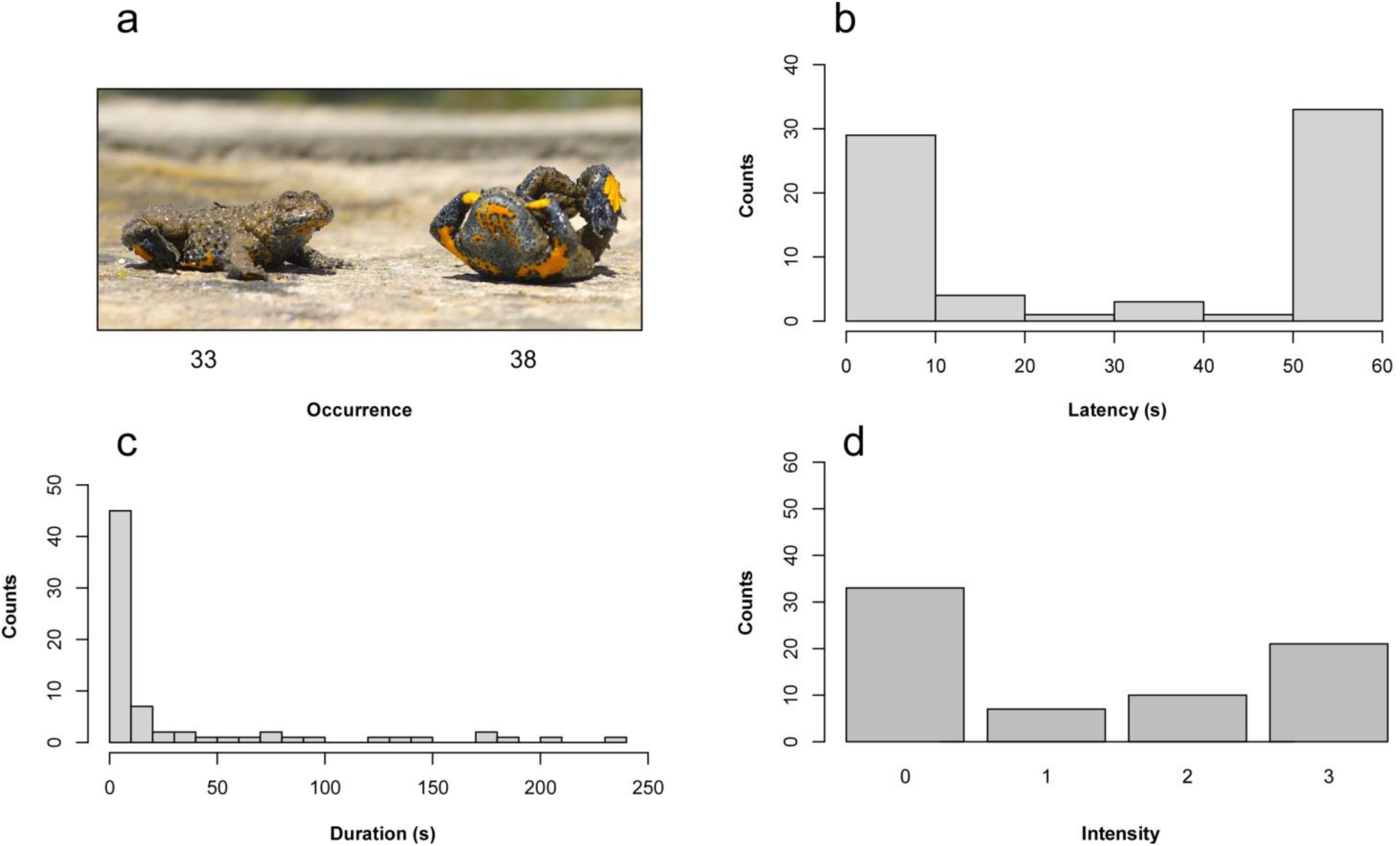
Representation of the inter-individual variation in the *unken-reflex* behaviour in the 71 tested *B. pachypus* individuals: a) image representing a non-responding individual (left) and a high-responding individual (right), with relative occurrence (below); b) latency of the response; c) duration of the *unken-reflex* display; d) intensity of the response, expressed as a scale from 0 to 3, where 0 is no response and 3 is the most intense reaction. Data refer to the first trial.

## Discussion

Anti-predatory behavior can differ among individuals, although the extent of this variation has been poorly characterized in the context of deimatic behavior. In simulated attacks on the Apennine yellow-bellied toads, we found that deimatic behavior varies among individuals, and that this variation is repeatable. We observed about half of the toads reacting to a simulated attack by arching the body and exposing the aposematically-colored ventral side. The other half did not show any display, but rather moved away. These results provide novel quantitative evidence of inter-individual variation in deimatic behavior, and raise questions on the evolutionary processes shaping alternative defensive strategies within populations.

The inspection of the pattern of variation in the unken-reflex behaviour revealed inter-individual variation in all behavioral descriptors, i.e. latency, duration and intensity of the deimatic display (Figure 1). The latency and the intensity of the display showed a bimodal pattern, with the extreme phenotypes way more frequent than intermediate phenotypes. This pattern suggests that no display (flee) or alternatively rapid and intense deimatic displays perform better than intermediate behaviors. The duration of the display has a uniform distribution, despite the display lasted more than one minute for most of the individuals performing the *unken-reflex*. This pattern suggests best performance of long displays respect to short display in halting predator attacks. The best performance of extreme behaviours is confirmed by the significant correlation shown by these traits (Table 2), which allows to define two alternative antipredatory strategies in *B. pachypus*: individuals quickly doing unken-reflex, with high intensity and long duration of the display, versus individuals avoiding the unken-reflex but rather escaping. These results support the occurrence of a trade-off between speed and accuracy [41] and are in line with the hypothesis that intermediate deimatic phenotypes could produce ambiguous stimuli, failing in startling or halting the predator attack [1].

We found all the behavioural descriptors highly repeatable across multiple trials (Table 1). This evidence has two main implications. First, the inter-individual variation in deimatic behavior of *B. pachypus* is consistent through time. Evidence of consistent behavioral differences among individuals within species have been accumulating in the last years and had major implications on the study of animal personalities (reviewed in [42]). Recent studies on the katydid crickets *Acripeza reticulata* revealed within-species differences in deimatic display (linked to the sexual dimorphism), but these differences were not consistent across multiple trials [43]. Our results show that deimatic behavior can consistently vary among individuals within species, and could therefore be linked to other personality traits. The second implication of this result is that the tonic immobility test applied here is repeatable, and it can be employed to quantify consistent individual differences in deimatic behaviour across multiple contexts, which is a promising subject for future investigations on behavioural responses in *Bombina* toads.

The two antipredatory strategies of *B. pachypus* could reflect the way individual cope with environmental challenges, as described by the coping style theory [44,45]. Indeed, employing deimatic display or rather escaping can be linked to the reactive-proactive coping styles. Accordingly, the reactive pattern involves a highly parasympathetic reactivity, resulting in a conservation-withdrawal response, conditioned immobility, and low levels of aggression [44,45,46]. This pattern fits with the rapid employing of deimatic display observed in *B. pachypus*. Conversely, individuals adopting a proactive coping style have a higher sympathetic activity and are characterized by active avoidance behaviour (commonly fight-or-flee response) and high levels of aggression. This pattern fits with the escaping behaviour observed in the toads not employing deimatic display. Further behavioral, physiological and genetic investigations (i.e. gene expression profiles) are mandatory to define deimatic phenotypes according to the coping style theory.

Overall, these results show that the *unken-reflex* is not a stereotyped deimatic behavior in *Bombina* toads. This evidence provides interesting insights on the evolutionary pathways to deimatism in *Bombina*, as well as on the mechanisms maintaining alternative antipredatory strategies within populations. Such consistent variation also lends support to the hypothesis that those behavioural patterns traditionally considered as rigid and stereotyped could have their evolutionary origin in phenotypic plasticity and learning processes [47]. The effectiveness of deimatic display, when chemical defences are also involved – as in the case of *Bombina* toads – does not require the predators learn the signal from the focal prey. Yet, predators can learn to ignore the deimatic component, but not the chemical defence [23]. In this frame, possessing a defensive ‘repertoire’ could provide populations with the opportunity to get away with a wider range of predators, including predators who learned to ignore the deimatic component. Therefore, the presence of such behavioural polymorphism would be adaptive at population level. However, more data are needed on extent of defensive repertories within populations of other deimatic species, either with and without further defenses. Moreover, further investigations should be addressed to evaluate the link between the observed variation and other variables, like personality traits, predator-exogenous stimuli, and the association between phenotypic and genetic variation.

## Acknowledgments

We are grateful to Giada Spadavecchia and Giacomo Grignani for their help in toad housing and feeding.

## Funding

This study was supported by the Aspromonte National Park.

## Competing interests

We declare that we have no competing interests.

## Notes

### Competing Interest Statement

The authors have declared no competing interest.

