## Supplementary material for "Shock or jump: deimatic behaviour is repeatable and polymorphic in a yellow-bellied toad"

### Supplementary methods

*Study species*

The Apennine yellow-bellied toad *B. pachypus* is an amphibian endemic to the Italian peninsula. It is a diurnal, thermophilic toad, active from April to October, and breeding in small seasonal ponds, from 100 up to 1600 masl. According to the altitude and local climate conditions, it starts reproducing from May to July; females produce up to 60 eggs and tadpoles usually metamorphose after one month; juveniles reach sexual maturity after two years. Adults' diet is composed by small invertebrates. This species is characterized by a dorsal cryptic habitus, and a ventral yellow-black aposematic coloration, which is exposed to the potential predator through arching the body, suddenly releasing toxins from skin glands (i.e. the *unken-reflex* deimatic behavior). The main predators of *B. pachypus* are birds and larger aquatic animals like grass snakes (*Natrix natrix*). Once abundant through the Italian peninsula, populations have been declining because of a combination of habitat alteration, climate change and disease (Lanza et al., 2007).

*Sampling and housing details*

Males were identified by the presence of forelimb nuptial pads. All the collected individuals were transported within three days to the experimental facilities, and housed individually in cages (25 cm × 25 cm × 25 cm). Toads were provided with a water tank and an oak wood as shelter. They were fed ad libitum with mealworm beetle larvae (*Tenebrio molitor*) and crickets (*Acheta domestica*). The cages were placed in a temperature-controlled room set to 24/25°C (monitored using a Hobo MX2201 temperature data-logger), 60%-80% of humidity, and natural photoperiod (Staniszewski, 1995).

Before any test started, we ascertained that the day-night activity rhythms were normalized (higher activity at night-time) by hourly visual check for two weeks; observations included reactivity to food and calling activity at night. No adverse effects on the overall health toad conditions were observed during the procedures. Finally, we checked for the occurrence of Chytrid fungus (*Batrachochytrium dendrobatidis*) infection in all the individuals: skin swabs were collected from all the individuals, and the molecular diagnostic essay was performed following the procedures described in Zampiglia et al. (2013). All the individual tested negative and could attend the behavioural tests.

Sampling procedures were performed under the approval of the Italian Ministry of Ecological Transition and the Italian National Institute for Environmental Protection and Research (ISPRA; permit number: 20824, 18-03-2020). Permission to temporarily house amphibians was granted by Local Health and Veterinary Centre, with license code 050VT427. All handling procedures outlined in the present study followed the Ethical Committee of the University of Tuscia for the use of live animals.
